# STIL-a novel link in Shh and Wnt signaling, endowing oncogenic and stem like attributes to colorectal cancer

**DOI:** 10.1101/2020.07.30.226654

**Authors:** Tapas Pradhan, Vikas Kumar, H Evangeline Surya, R Krishna, Samu John, VT Jissa, S Anjana, K Chandramohan, S Asha Nair

**Affiliations:** Cancer Research Program 4, Rajiv Gandhi Centre for Biotechnology, Trivandrum, Kerala, India; Department of Surgical Oncology, Regional Cancer Centre, Trivandrum, Kerala, India; Department of Statistics, Sree Chithra Tirunal Institute of Medical Science and Research, Trivandrum, Kerala, India

**Keywords:** Colorectal cancer, STIL, Hedgehog Signaling, Cancer stem cell, Drug resistance, β-catenin, Prognosis

## Abstract

Discovery of potent gene regulating tumorigenesis and drug resistance is of high clinical importance. STIL is an oncogene, however its molecular insights and role in colorectal oncogenesis are unknown. In this study we have explored role of STIL in tumorigenesis and studied its molecular targets in colorectal cancer (CRC). STIL silencing reduced proliferation and tumor growth in CRC. Further, STIL was found to regulate stemness markers CD133 & CD44 and drug resistant markers Thymidylate synthase, ABCB1 & ABCG2 both in in-vitro and in-vivo CRC models. In addition, over expression of STIL mRNA was found to be associated with reduced disease free survival in CRC cases. To our surprise we observed an Shh independent regulation of stemness and drug resistant genes mediated by STIL. Interestingly, we found an Shh independent regulation of β-catenin mediated by STIL via p-AKT, which partially answers Shh independent regulatory mechanism of CSC markers by STIL. Our study suggest an instrumental role of STIL in molecular manifestation of CRC and progression.

## 1 Introduction

*SCL/TAL1* interrupting locus (STIL) gene is a crucial factor in centriole biogenesis and dysfunction of this gene has been associated with abnormal brain development leading to microcephaly (1). Being a 1288 amino acid cytoplasmic protein, STIL functions as a cell cycle-regulatory protein specifically recruited at the mitotic centrosome to promote the duplication of centrioles in dividing cells. STIL has been shown to interact with CDK1, PLK4, SAS-6,which are crucial in centriole duplication (2) and thus proven to have a crucial role in cell division (3)In lung cancer, up-regulation of STIL has been reported to have significant effect on tumor mitotic activity(4). STIL also has been reported to be over expressed in pancreatic ductal cell carcinoma(5) and altered in leukemia (6). Further, STIL overexpression has been reported tocause chromosomal instability in cancer cells (7). Being a cytoplasmic protein, STIL has also been studied as an integral part of hedgehog signalling (Hh) cascade. The C terminus of STIL can interact with conserved components of the Hh signalling such as suppressor-of-fused homolog (SUFU) and GLI1. Interaction of STIL with SUFU inhibits the repressor function of SUFU towards GLI1 resulting in activation of Hh-Gli1 cascades. Knockdown of STIL has been shown to increase nuclear accumulation of SUFU with GLI1 and repression of GLI1 mediated transcriptional activity (5). STIL^-/-^ mouse embryos have been shown to have reduced expression of in PTCH1 and Gli1 and has been reported to lack primary cilia, a structure present in almost all cell types mediating Hh signaling (8,9). During the process of carcinogenesis, increased expression of STIL promotes the transcriptional activity of GLI1 leading to increased transcription of GLI1 targets that promote sustained proliferation, cell death resistance, stemness, angiogenesis, and genomic instability, which are the hallmarks of cancer (10). Therefore, increase in STIL expression likely represents a crucial step toward cancer progression. STIL depletion has been shown to enhance DNA double strand breaks caused by DNA damaging agents in ovarian cancer (11). Another report has shown STIL to play a sonic Hh dependent role in drug susceptibility in PC12 cells(12). Further Hh signalling has been studied for its role in maintenance of CSC, metastasis and disease recurrence in colorectal cancer (CRC) (13). GLI1 mediated Hh signalling has been found to have important role in CRC cell survival upon therapeutic insults (14). However, there have been hardly any studies deciphering the role of STIL in CRC tumorigenesis and drug resistance. Studies have shown regulation of ABCB1 and ABCG2 efflux pumps to be Hh dependent, which suggest that the Hh pathway could be a potent target to overcome drug resistance and surge chemotherapeutic response (15). Nevertheless, tumor fate against therapies is governed by various cues from tumor microenvironment and thus a complex network of inter and intra cellular signaling decides therapy response. There has been reports suggesting instrumental role of Wnt-Shh crosstalk in basal cell carcinoma (16) and gastric cancer (17). In addition, crosstalk between Wnt and Shh has been shown to be contributing towards CRC progression (18), however core pathway components mediating crosstalk still remain unexplored. Further studies to explore the mediators of this crosstalk would provide an in depth mechanistic prospective on tumor development and therapy resistance in CRC. In this study we have explored the multifaceted role of STIL in CRC and also deciphered its role in mediating cross talk between Shh and Wnt signaling.

## 2 Methods

### 2.1 Patients’ Samples

Biopsies were collected from CRC patients undergoing curative surgery between 2014 and 2017 at the Regional Cancer Center, Trivandrum after human ethics committee approval and sanction from Institutional Review Board*. All subjects gave written informed consent in accordance with the Declaration of Helsinki*. Patient’s details and clinical information were collected from medical records of the same institution.

### 2.2 Cell culture

HCT116, HT-29 & HEK293T cells (ATCC, USA) were cultured with Dulbecco’s Eagle’s Medium (DMEM) (Invitrogen, USA) supplemented with 10% FBS (Invitrogen, USA) at 37°C and 5 % CO2 in a humidified incubator (Thermo scientific, USA).

### 2.3 Chemical inhibition of Shh signaling

(4-Benzyl-piperazin-1-yl)-(3,5-dimethyl-1-phenyl-1H-pyrazol-4-ylmethylene)-amine (SANT1), a chemical inhibitor of SMO receptor was used at a concentration of 30nM for 24 hours to inhibit Shh signaling in HT29 cells. Post treatment cells were processed for RNA and protein isolation.

### 2.4 RNA Isolation

Trizol reagent (Invitrogen,USA) was used to isolate total RNA from around 3×10^6^ adherent cells or 50 mg tissues following manufacturer’s protocol. RNA quality and quantification was carried out on Nanodrop 1000 (ThermoScientific,USA) after assessing its quality using gel electrophoresis.

### 2.5 Real-time q-PCR

1μg of RNA was used for conversion of cDNA from each sample using PrimerScript cDNA conversion kit (TAKARA, Japan), following manufacturers’ protocol. Quantitative real-time PCR was performed using SYBR-Green based fluorescence detection kit (TAKARA, Japan) and HT9700 detection system (AB, Life science, USA). 25 ng of cDNA was used as template for each reaction. Analyzed genes and the primers used are shown in **Supplementary Table 1.** PCR data were analysed using Data assist software (AB, Life science, USA).

### 2.6 Lentiviral mediated gene silencing

HEK293T cells (3×10^5^) were seeded in a 6 well plate (Nunc,Thermo,USA) with DMEM media (Invitrogen,USA) containing 10% FBS (Invitrogen,USA) and incubated till it reached 60% confluency.DNA constructs were mixed in a fixed ratio in an vial containing 75 μl OptiMEM media (Invitrogen,USA) as shRNA(0.75 μg/well), pREV(0.5 μg/well), pMDL(0.18 μg/well), pVSVG(0.26 μg/well) (Packaging plasmids were gifted from Vinay Tergaonkar, Department of Biochmeistry,Yong Loo Lin School of Medicine, NUS). Details of shRNA employed are given in **Supplementary Table 2**. To the mixture of constructs 2mg/ml polyethylenimine (PEI)(Sigma-aldrich,USA) was added followed by brisk vortex for 10 sec and incubated for 30 min at RT. During incubation HEK293T cells were rinsed with OptiMEM media and 688μl of OptiMEM was added to each well. After incubation 75 μl DNA/PEI complex was added to cells drop wise followed by gently shaking and cells were kept in incubator overnight. Fresh 1.5ml DMEM media with 10% FBS was added to the cells after overnight incubation and media was collected every 24 hrs from cells and replenished by fresh media for 3 consecutive days. Collected media was filtered using 0.45μm PES membrane (Minisart, Sartorius, Germany) to remove extracellular debris. Filtered media containing lentiviral particles were centrifuged at 24000rpm at 20°C for 2 hrs in 90Ti rotor in ultra-centrifuge (Beckman Coulter Optima TM L-100K). Viral particle pellet was obtained after discarding supernatant post centrifugation and suspended in 100 μl OptiMEM media for overnight at 4°C.The collected lentivirus suspension was used for transfection or stored for later use at −80°C. 3× 10^5^ number of HT29 and HCT116 cells were seeded in 6 well plate in DMEM media containing 10% FBS and incubated till it reached 70% confluency. After reaching desired confluency, cells were infected with 100μl of lentiviral particles suspension along with 10 μg/ml polybrene (Sigma-aldrich,USA) 1.5ml of fresh DMEM media containing 10% FBS without antibiotics and incubated for minimum of 24hrs. Cells were selected using puromycin (Sigma-aldrich,USA) antibiotic with a concentration of 2μg/ml and 7.5μg/ml for HCT116 and HT29 respectively for 3 weeks for the development of stable transfected cells. Gene knock down was evaluated using real-time qPCR and western blot.

### 2.7 Cell Proliferation assay

MTT-3-(4, 5-Dimethylthiazol-2-yl)-2, 5-Diphenyltetrazolium Bromide) assay was used to determine cell survival. 100 μl of culture medium containing 1×10^6^ cell were seeded into a 96 well plate and incubated for 24hrs. After that 0.5mg/ml of MTT was added to each well and incubated for 3hrs in dark at 37°C. Culture medium from the wells was aspirated and 100μL of iso-propanol was added to dissolve the formazan crystals formed inside cells. Absorbance was measured at 570 nm in a microtiter plate reader. A graph was plotted with concentration on X axis and absorbance on y axis.

### 2.8 Cell cycle analysis

Cells were harvested after trypsinization by centrifugation at 1200g for 5 min at 4°C. 300μl of ice-cold PBS was used for the suspension of cells and 700μl of 70% ethanol was added drop wise with slight vortex followed by incubation for 1 hr in ice. Post incubation cells were centrifuged at 1200g for 5 min at 4°C and cells pellet was suspended in 1X PBS. Cells were treated with RNase A (Sigma-aldrich, USA) at a concentration of 100μg/ml for 1 hr at 37°C in a heat block (Thermo mixer, Eppendorf, Germany). After incubation 10μg/ml of propidium iodide was added in dark for 15 min. Cells were strained using 45μm cell strainer and collected in FACS tube. Tubes were kept in ice till flow analysis.

### 2.9 Immunophenotyping

Cells were harvested after trypsinization by centrifugation at 400g for 5 min and washed with 1ml HBSS. 1×10^6^ cells were used for antibody staining according to manufacturer’s protocol. Details of antibodies used are provided in **Supplementary Table 3.** After antibody staining, cells were washed with ice cold HBSS and suspended in HBSS containing 2% FBS. Cells were kept on ice till flow cytometry analysis was carried out using BD FACS ARIA II (BD Bioscience, USA).

### 2.10 Side population analysis

Side population assay was performed according to the protocol developed by Goodell (1996) with slight modifications. Harvested cells were spinned at 400g for 5 min and pellet was suspended at 1×10^6^ cells per ml in pre-warmed DMEM (supplemented with 2% serum) media (Gibco, Invitrogen) in two tubes. Hoechst 33342 (Sigma-aldrich) was added to a final concentration of 5 μg/ml at dark in both tubes. In control tube 100μm verapamil was added to cells prior to addition of 5 μg/ml Hoechst. Cells were mixed well and placed in 37°C dry bath(Thermomixer, Eppendorf,Germany) with shaking at 500rpm. After incubation cells were spinned down at 4°C and re-suspended in cold HBSS (with 2% serum). Cells were suspended in ice cold HBSS (2% serum) containing 2 μg/ml Propidium iodide (Sigma-aldrich,USA) for dead cell discrimination. Cells were immediately taken for flow cytometry analysis (BD FACS Aria II) and results were analyzed using FACS Diva software.

### 2.11 Apoptosis assay

Cells were seeded in 6 well plate and incubated for 24hr.5-Flurouracil was added to wells at a sub lethal concentration keeping DMSO treatment as control for 24hr.Cells were washed with 1X PBS followed by trypsinization and pelleted by centrifugation at 400g for 3 min at 4°C.

Cells 100μl were suspended in 1X binding buffer and 5μl of Annexin reagent and incubated in dark for 15 min at RT. After incubation 5μl propidium iodide was added to cells and kept for 5 min. Finally 400μl of 1X binding buffer was added and cells were filtered using 40μm strainer and analyzed by flow cytometry using BD FACS Aria II (BD Bioscience,USA).

### 2.12 Protein isolation

The cells were washed with 1ml 1x PBS. 1ml 1x PBS was added and the cells were scraped out using a cell scraper. The cells were collected into an eppendorf and centrifuged at 5000 rpm for 5 min at 4° C. The supernatant was discarded and 50-200μl of RIPA buffer was added based on the pellet size. Pho and PI (10μl/1ml buffer) were added. The pellet was suspended well and vortexed. Mixed in thermomixer at 4° C for 1.40 hours at 1400rpm and centrifuged at 14000rpm for 15 minutes at 4° C. The supernatant was collected and stored at −80° C.

### 2.13 Nuclear and Cytoplasmic protein fraction isolation

Approximately 1 x 10^7^ cells were washed with 1x PBS and harvested by centrifugation at 400g for 5 min. The pellet was resuspended in 5 pellet volume of CE buffer (1X solution composed of 10 mM HEPES, 60 mM KCl, 1 mM EDTA, 0.075% (v/v) NP-40, 1mM DTT and 1 mM PMSF, pH 7.6),incubated on ice for 3 minutes and spun at 1500 rpm for 4 minutes. The supernatant (cytoplasmic extract) was removed into a clean tube and the pellet was washed with 100 μL of CE buffer without NP-40 at 1500 rpm for 4 minutes. One pellet volume of NE buffer (1X solution composed of 20 mM Tris Cl, 420 mM NaCl, 1.5 mM MgCl2, 0.2 mM EDTA, 1mM PMSF and 25% (v/v) glycerol, pH 8.0)was added to the pellet, salt concentration adjusted to 400 mM using 5 M NaCl (~35 μL), and an additional pellet volume of NE buffer added to it and resuspended by vortexing. It was incubated on ice for 10 minutes with periodic vortexing. Both CE and NE were then spun at maximum speed (14,000 rpm) for 10 minutes. The supernatants from all the tubes were then transferred separately to clean tubes and stored at −80 °C.

### 2.14 Western blot

After SDS-PAGE separation of proteins transfer sandwich was prepared with PVDF membrane and placed in transfer buffer. PVDF membrane (Amersham) was activated with methanol for 15 sec prior to use. The blot was allowed to run for 120 min at 100 volts. After the transfer of proteins, membrane was blocked using 5% skimmed milk powder as blocking buffer for 1hour at RT. Then membrane was washed with 1X TBST buffer and incubated with primary antibody for overnight (Details of antibody in **Supplementary Table 3**). Again membrane was washed in 1X TBST buffer followed by secondary antibody (1:5000dilution) treatment, then incubated for 1hr at RT. After incubation membrane was washed with 1X TBST buffer 3 times and detected with ECL reagent (TAKARA, Japan) using Versa Doc (BD Bioscience, USA) or X-ray film.

### 2.15 Development of tumor Xenograft

Each experimental groups had five 8 weeks old male NOD/SCID mice, which were housed in a specific pathogen-free environment. Four million cells/ mice of each control shRNA & STIL-shRNA HT-29 cells were suspended in PBS and injected into flanks subcutaneously. Then mice were monitored every two days for tumor growth and health. Once visible tumor appeared, its volume was measured using callipers every two days for obtaining rate of tumor growth in both groups. Tumor volume was calculated using the formula V(mm^3^) = (Largest length) × (Shortest length)^2^ /1(19). At the end mice were sacrificed using carbon dioxide inhalation and tumor along with lungs and liver tissues were harvested and stored in RNA Later and Formalin for molecular and histological analysis respectively.

### 2.16 H & E staining

Haematoxylin and Eosin (Merck-millipore, USA) staining was be performed according to manufacturer’s protocol for pathological examination of various tissues obtained after surgery.

### 2.17 Immunohistochemistry

Tissues received post-surgery were fixed with 4% paraformaldehyde (Sigma-aldrich, USA) at 4°C for 16 hours. They were then processed with alcohol gradient and xylene followed by embedding with paraffin to make paraffin blocks. 5 μm thick sections were mounted on Starfrost glass slides (Leica, Germany) and were deparaffinised in xylene followed by rehydration with alcohol gradients. Sections were blocked for endogenous peroxidases and antigen retrieval was performed using 10 mM citrate buffer (pH 6.0). After retrieval BSA (3%) blocking was done for 30 min at RT followed by primary antibody staining for overnight. Primary antibodies details are given in **Supplementary Table 3**. After incubation antigen-antibody binding was detected using ABC kit (Vecta stain, USA). DAB substrate (Sigma-Aldrich, USA) was used as a chromogen and hematoxylin (Merck Millipore, Germany) was used as a counter stain. Sections without incubation of primary antibody served as negative control. Semi quantitative analysis was done by counting three independent microscopic fields (n=100-200 cells/field) for staining of specific antigen using upright microscope (Leica DM1000, Germany). Tissue array (US Biomax, USA) for STIL expression was also performed according to above protocol. Tissue array demographics has been detailed in **Supplementary Table 4**.

### 2.18 Cancer Datasets analysis

STIL gene was used as an input in Oncomine (www.oncomine.org/resource/) online free cancer database to obtain gene expression pattern across multiple colorectal cancer transcriptomics studies globally and survival curves for STIL expressing patients were obtained from C-Bioportal (www.cbioportal.org) online tool.

### 2.19 Statistical Analysis

All statistical analysis was performed using SPSS version 25 and GraphPad Prism 5. Unpaired t-test was performed for mean comparison and p value determination for flow cytometry and immunohistochemistry data.. Real-time PCR data analysis was carried out using one sample t test to compare determined log2 fold change and determined p value among samples. Kaplan-Meier plots were obtained from C-Bioportal cancer genomic datasets. Multivariate analysis was performed using binary logistic regression model and p-value and ODDS ratio were derived.

## 3 Results

### 3.1 Over expression of STIL gene in CRC correlates with lower disease free survival in patients

We used Oncomine datasets to analyze differential mRNA expression of STIL between normal and cancerous colon tissues from three different study (Hong, Notterman & Kaiser), which revealed a significant enrichment of STIL gene in cancer tissues (**Figure 1A**). Further validation in our cohort CRC using real-time q-PCR showed a 2.1 fold high expression in tumor compared to matched normal tissues (p = 0.001) (**Figure 1B**). IHC of STIL protein in cancer tissues showed a very high cytoplasm specific staining compared to normal tissue (**Figure 1C**). To assess the role of STIL enrichment in colon cancer with disease prognosis, we used c-Bioportal cancer database. Kaplan-Meier plots showed no significant differences for 5-year survival among STIL high and low expression groups, however STIL high expression group showed a very low disease free survival rate among CRC patients with a p value 0.001 and 0.0007 in two different studies (**Figure 1D**). Put together, these results suggest that over expression of STIL in CRC could have crucial unexplored role in tumor development and disease progression.

**Figure 1.**
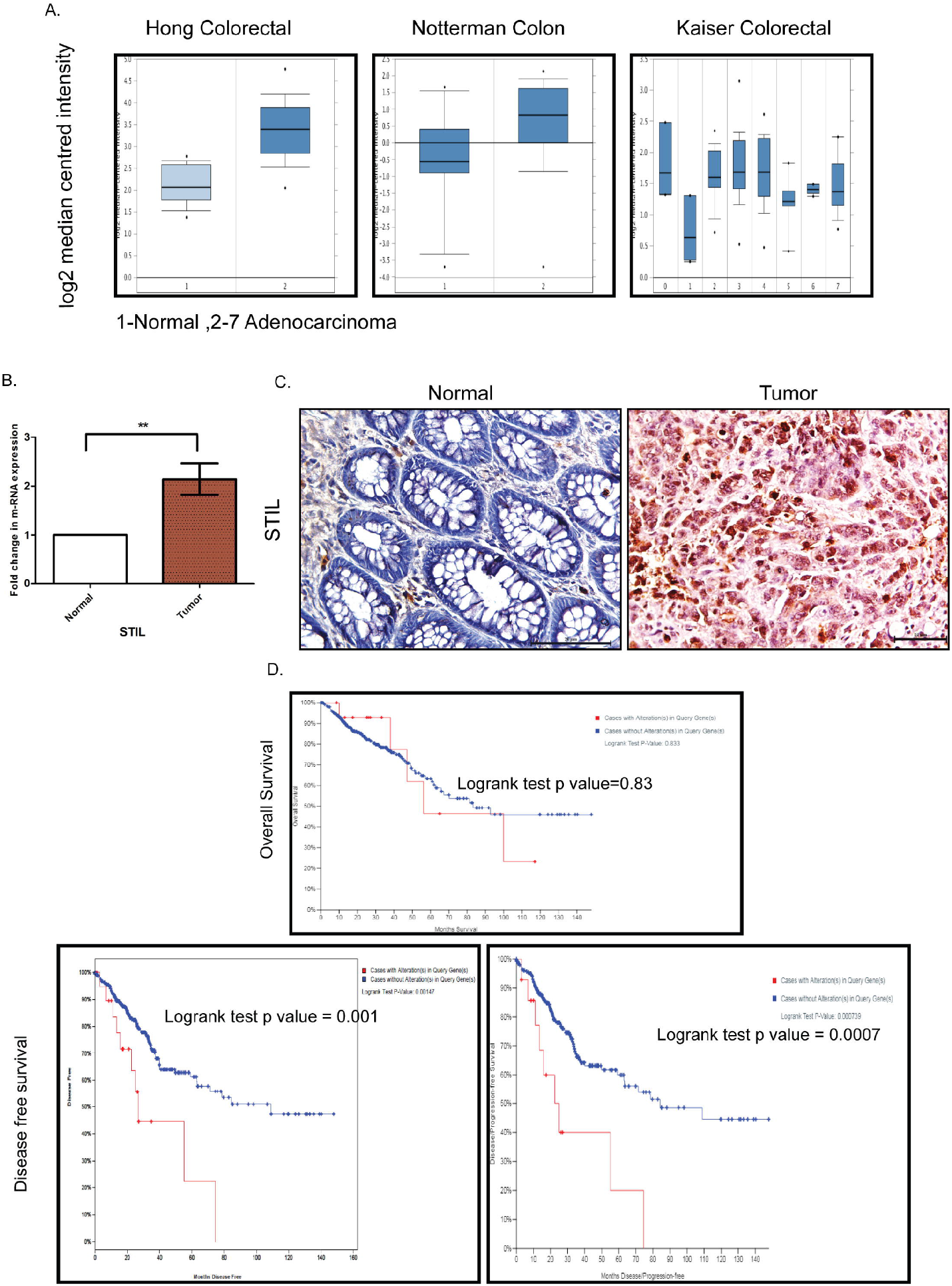
Expression of STIL gene in CRC and its role in prognosis. (A) STIL m-RNA expression in normal colon and tumor obtained from Oncomine datasets. (**B & C**) Differential expression of STIL gene mRNA (N=22) and protein in CRC.(**D**) Kaplan-Meier plot showing association of high mRNA expression with patients’ survival and disease free survival in CRC cases. One Sample t-test and log rank test results showing p value ≥0.05, ≥0.01, are represented by *,** respectively, Scale bar is 20μm.

### 3.2 High STIL protein expression found to have association with early tumor stage in rectal cancer

To evaluate STIL protein expression in different tumor stages and grades of CRC, we performed IHC for STIL in rectal cancer tissue array (Array details given in **Supplementary Table 4**). STIL was found to have intense cytoplasmic staining in cancer tissues compared to normal (p value 0.001). All tumor stages were found positively stained with a high IHC score associated with early stage (T1-T2) compared to late stage (T3-T4) tumors (**Figure 2A & Table 1**). Both high and low grade tumor tissues showed moderate to strong STIL staining except signet-ring carcinoma (**Figure 2B**). Chi-square analysis revealed that females (69.4%) and early stage tumor (73.3%) correlated well with higher expression of STIL with a p value of 0.04 & 0.03 respectively (**Table 1**). However, gender adjusted binary regression analysis revealed that higher tumor stage has 71% lesser chance of high expression of STIL compared to early stage (p value0.02) and earlier association of female gender with STIL expression was influenced by tumor stage. In addition, late disease stage was found to have 60% lesser chance of STIL expression compared to early stage (p value 0.07) (**Table 2**). In addition, role of STIL in cancer cell migration was evaluated using scratch wound healing assay, which showed no significant difference between control and STIL silenced HCT116 cells (**Supplementary Figure 7**). In addition, we also analyzed lungs and liver tissues for presence of secondary tumor in control and STIL silenced xenograft mice using Ep-CAM protein expression and found no tumor deposits (**Supplementary Figure 8**). This further suggest that STIL may not play a role in cancer cell migration and thus metastasis in concordance to tissue array observation. Put together, this study suggests that STIL may not play a significant role in invasion of tumor, however it could have crucial role in early tumorigenesis.

**Figure 2.**
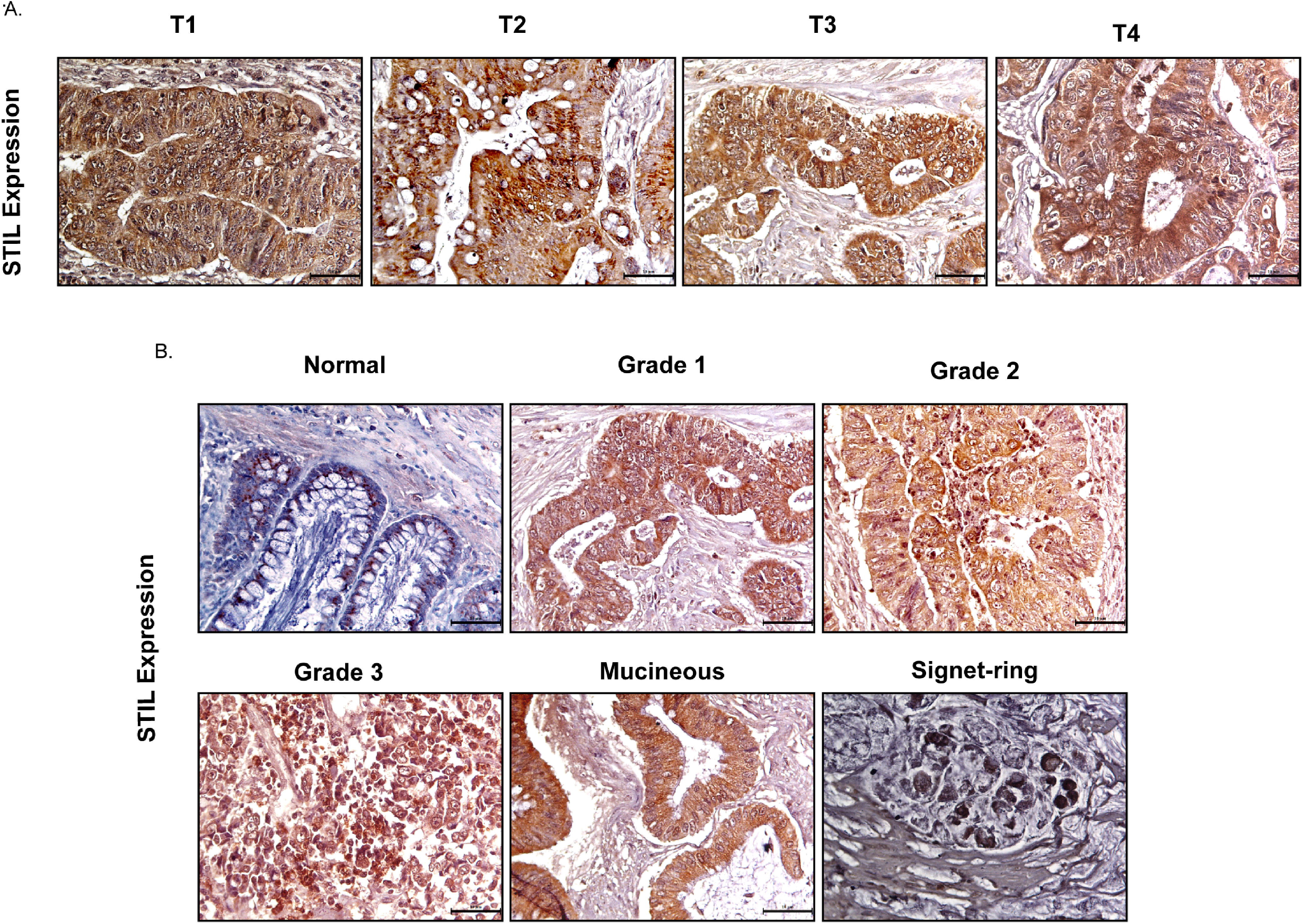
IHC of rectal cancer tissue array showing expression of STIL protein. (**A & B**) Representative images showing expression of STIL protein in different tumor stages and grades.

**Table 1.** Chi-square test showing association of clinical and pathological parameters with STIL protein expression.

|  |  | IHC score |  | Total % | P- value |
| --- | --- | --- | --- | --- | --- |
|  |  | Low% | High% |  |  |
| **Gender** | Female | (11) 30.56 | (25) 69.44 | (36) 100 | 0.04 |
|  | Male | (21) 53.85 | (18) 46.15 | (39) 100 |  |
| **Age (Years)** | ≤55 | (17) 45.95 | (20) 54.05 | (37) 100 | 0.57 |
|  | >55 | (15) 39.57 | (23) 60.53 | (38) 100 |  |
| **Pathology** | Normal | (3) 60 | (2) 40 | (5) 100 | 0.001 |
|  | Carcinoma | (21) 33.87 | (41) 66.13 | (62) 100 |  |
|  | Others | (8) 100 | 0 | (8) 100 |  |
| **Tumor Stage** | ≤T2 | (8) 26.67 | (22) 73.33 | (30) 100 | 0.03 |
|  | >T2 | (21) 52.50 | (19) 47.50 | (40) 100 |  |
| **Disease Stage** | I | (8) 28.57 | (20) 71.43 | (28) 100 | 0.07 |
|  | II & III | (21) 50 | (21) 50 | (22) 100 |  |

**Table 2.**
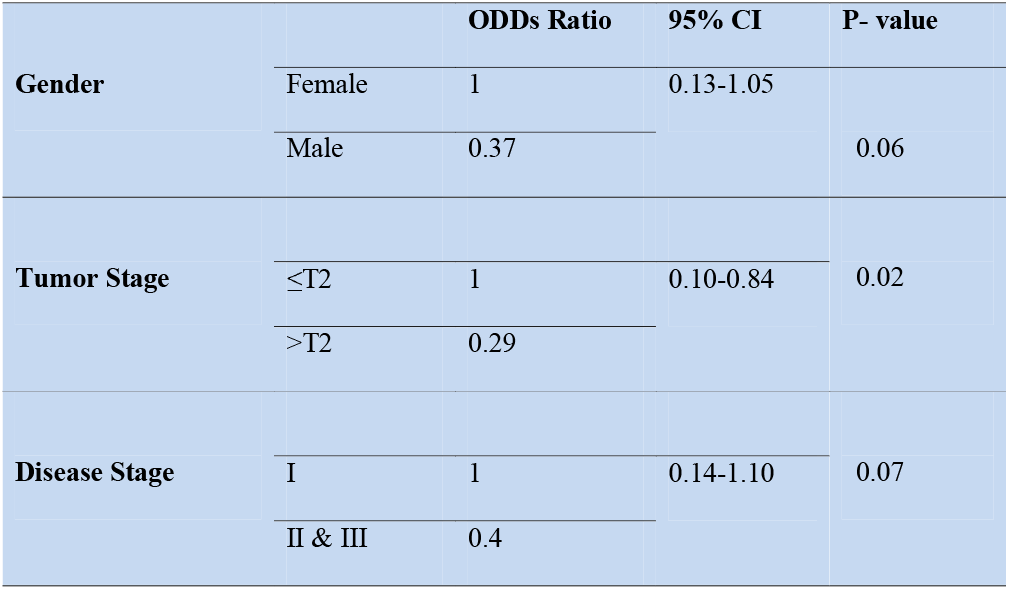
Binary regression analysis showing gender adjusted association of tumor and disease stage with STIL expression.

### 3.3 STIL silencing reduced cell proliferation and tumor growth in CRC

To understand the role of STIL in proliferation and tumor growth of CRC, we stably silenced STIL gene in HT-29 CRC cells using shRNA (**Figure 3A & B**). Cell survival was evaluated by MTT viability assay, where STIL silencing reduced the proliferation by 34% compared to control shRNA cells (p=0.05)(**Figure 3C**). STIL has been well studied for its role in cell cycle regulation, as it plays a crucial role in centrosome assembly. Thus, we analyzed cell cycle pattern, which revealed a twofold accumulation of cells in G2/M phase in STIL silenced cells (**Figure 3D**). These results led us to investigate the tumorigenic potential of STIL *in-vivo*, for which we developed a tumor xenograft using STIL silenced HT-29 cells and compared it with control shRNA xenograft. We observed a higher tumor volume in control group which became more significant with time compared to STIL silenced tumor (**Figure 3E & F**). Dry weight of control tumor was found to be 3 fold more than STIL silenced tumor (p=0.001) (**Figure 3G**). Further, we analyzed nuclear expression of the proliferation marker Ki-67 in tumor xenograft tissue and found a significantly reduced expression in STIL silenced group compared to control (p=0.01) (**Figure 3H**). These results confirm a significant role of STIL in regulating tumor proliferation and growth in CRC.

**Figure 3.**
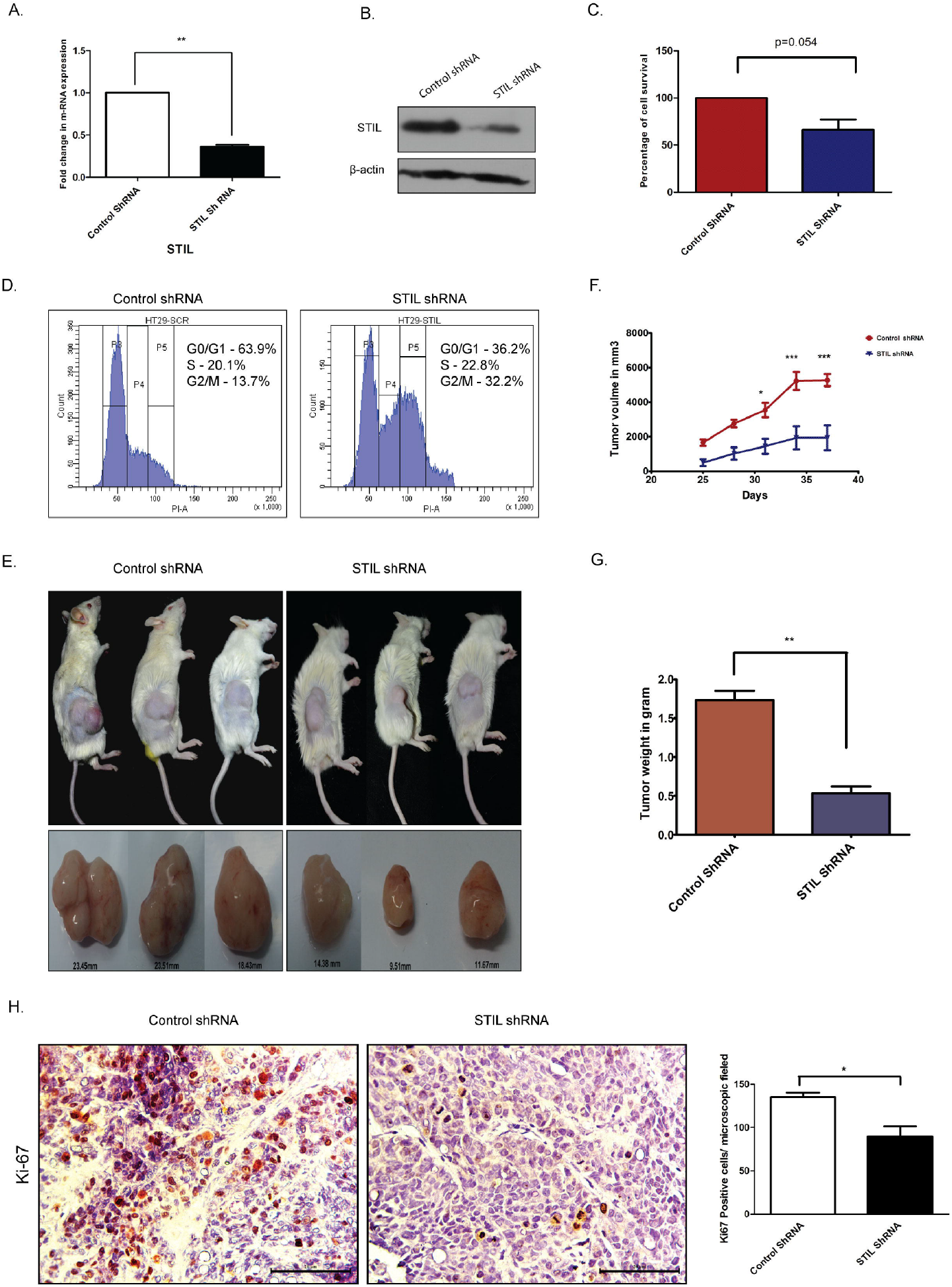
Role of STIL in CRC tumorigenesis. (**A**) Bar graph showing fold change in STIL mRNA expression upon STIL silencing. (**B**) Immunoblot validating STIL silencing at protein level. (**C**) Bar graph showing cell proliferation of HT-29 cells upon STIL silencing. (**D**) Histogram showing effect of STIL silencing on cell cycle. (**E & F**) Representative images of tumor xenograft and graph showing growth of tumor upon time (n=3). (**G**) Bar graph showing mean dry weight of tumor in both groups (n=3). (**H**) IHC showing nuclear expression and quantification of Ki-67 protein in xenograft derived tumor tissues. Unpaired t-test results showing p value ≥0.05, ≥0.01, ≥0.001 are represented by *,**,*** respectively, Scale bar is 20μm.

### 3.4 STIL regulate cancer stem cell associated genes expression in CRC independent of Shh signaling

We have observed STIL over expression to be associated with lower disease free survival in CRC patients from our initial results. Disease recurrence has mostly been contributed by residual cells, which are drug resistant in nature. These cells are known to have cancer stem cell (CSC) properties including expression of established surface markers. However, role of STIL in CRC stem cells is still unknown. We assessed the expression of established CSC markers in STIL silenced cells and observed a significant reduction in CD133 (fold change 0.4, p=0.03) and CD44 (fold change0.7, p=0.02) expression in m-RNA and protein (**Figure 4A & B**). STIL is a known positive regulator of Shh signaling and thus, we analyzed expression of CSC markers CD133 & CD44 upon Shh inhibition by SANT1 treatment and found no significant reduction in their expression, suggesting a hedgehog independent regulation of CSC markers by STIL (**Figure 4C & D**). We further validated these findings in xenograft tissues using real-time q-PCR, which showed significant reduction in CD133 (fold change 0.41, p=0.002) and CD44 (fold change 0.77, p=0.05) m-RNA expression in STIL silenced tumor, which was confirmed by IHC (**Figure 4E-G**) respectively. Additionally, we also analyzed expression of stem cell maintenance factors OCT4 and NANOG in STIL silenced cells and found a reduction (fold change 0.69 & 0.78) compared to control respectively, however statistical significance was not observed (**Supplementary Figure 9**). Membranous expression of CD133 and CD44 are known to be characteristics of CSC, thus we also analyzed their surface expression using flow cytometry. In concordance with our gene expression data, a significant reduction in CD133 and CD44 positive cells were also observed upon STIL silencing (**Figure 4H**). These results confirm a STIL mediated regulation of CSC signatures associated with CRC.

**Figure 4.**
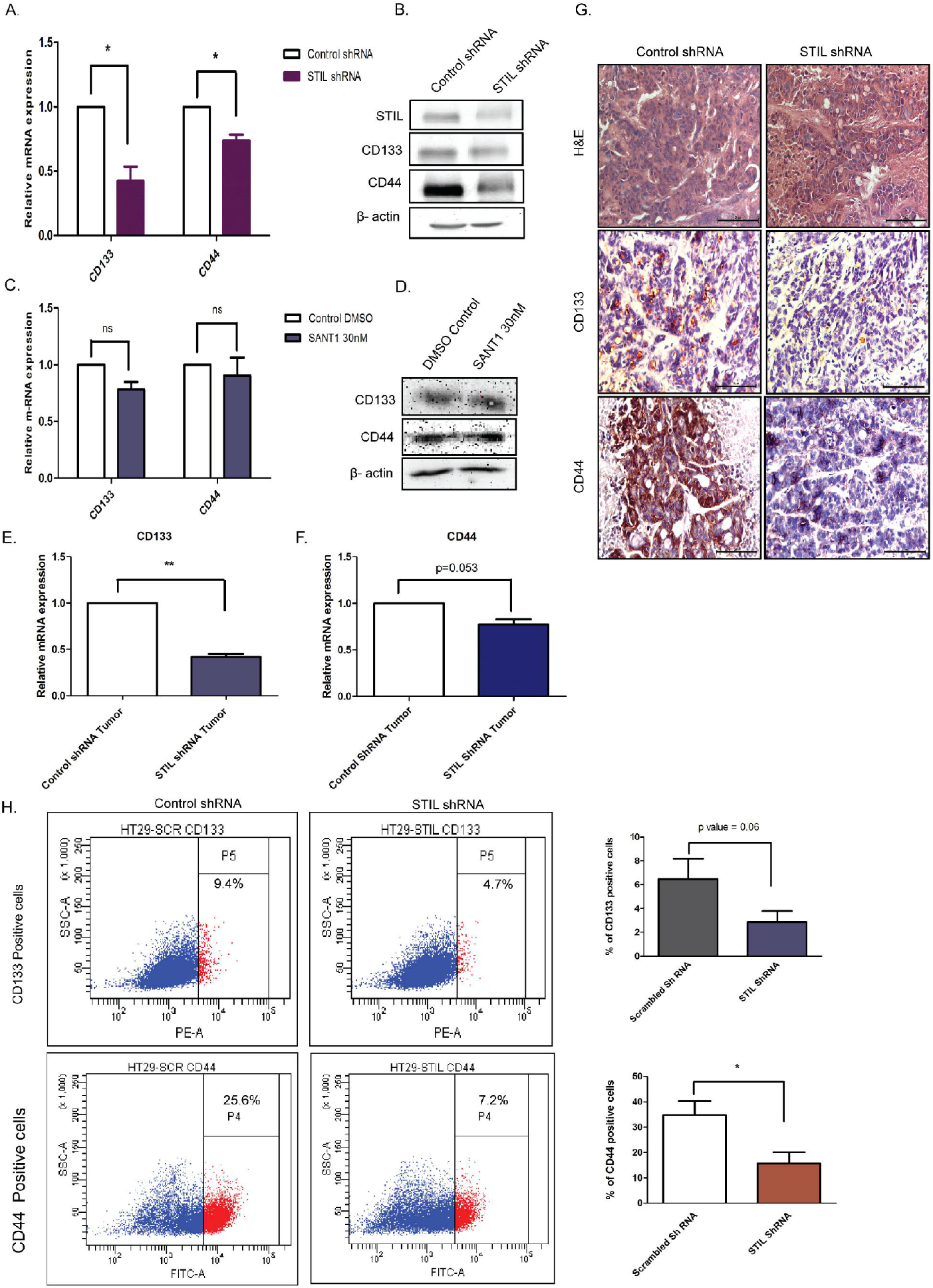
Effect of STIL silencing on CSC markers expression in CRC. (**A & B**) mRNA and protein level expression of CSC markers upon STIL silencing. (**C & D**) mRNA and protein level expression of CSC markers upon SANT1 treatment. (**E-G**) Showing CD133 & CD44 expression in xenograft tumors at mRNA and protein levels respectively. (**H**) Dot plot representing % of CD133&CD44 positive cells. Unpaired t-test results showing p value ≥0.05, ≥0.01, ≥0.001 are represented by *,**, *** respectively, Scale bar is 20μm.

### 3.5 STIL silencing reduced drug efflux activity via depletion of drug transporters proteins in CRC cells and sensitized cells for 5fu treatment

Another hallmark of CSC is the ability to efflux drugs via active expression of drug transporter proteins. Here, we performed side population (SP) assay which potentially identifies drug effluxing cells and CSC. STIL silencing resulted in reduced SP in HT-29 cells with mean % population of 0.7 compared to control with 3.7(p=0.051) (**Figure 4A & B**). Our data further revealed depletion of ABCB1 and ABCG2 proteins upon STIL silencing (**Figure 4F**). Moreover, ABCG2, which is known to be the most crucial transporter associated with stem cells and SP, was found to have reduced mRNA expression in both STIL silenced HT29 cells and xenograft tumor tissue (**Supplementary Figure 10A & B**). In concurrence with the mRNA data we also observed reduced ABCG2 protein expression in STIL silenced tumor tissues (**Figure 4G**). In addition STIL silencing was also found to deplete expression of thymidylate synthase (TS) enzyme, known for its role in 5-FU resistance in CRC (**Figure 4F**). However, we observed no reduction in ABCG2 and TS proteins expression upon Shh inhibition, revealing an Shh independent regulation of these proteins by STIL (**Figure 4E**). However, ABCB1 expression reduced significantly upon Shh inhibition (**Figure 4E**). Further, we analyzed effect of STIL silencing on 5-FU drug treatment in HT29 cells and observed enhanced cell death in STIL silenced cells compared to controls (**Figure 4C & D**), which could be the result of reduced drug efflux activity and thymidylate synthase expression. Put together, these data suggest an Shh independent regulatory role of STIL in CSC and drug resistance in CRC.

### 3.6 STIL regulate β-catenin expression via Akt independent of Shh signaling

STIL has been studied to be a positive regulator of hedgehog signaling, however STIL mediated hedgehog regulation in CRC remains unknown. To study its role in mediating regulation of hedgehog signaling in CRC, we screened for expression of hedgehog associated genes (SHH, PTCH2, SMO, SUFU & STIL) as well as effector genes (GLI1 & GLI2) after STIL silencing and treatment with SANT1. SANT1 treatment resulted in significant down regulation of the pathway associated genes as well as effectors molecules GLI1&2 (**Figure 6A**). Conversely, STIL repression did not show similar trend in gene expression except for GLI1 (**Figure 6B**). These observations suggest negative regulatory role of STIL on SUFU,SMO and SHH genes in CRC. Although GLI2 showed an increase in mRNA level upon STIL silencing, it showed a significant reduction at protein level upon STIL repression and Shh inhibition (**Figure 6C & D**). There are studies which advocate a reciprocal relationship between Wnt and Hedgehog signaling in CRC development (20). We further looked into probable cross talk between Shh and Wnt signaling and observed an interesting STIL mediated regulation of Wnt effector β-catenin, which was not observed upon Shh inhibition (**Figure 6C & D, Supplementary Figure 11**). Further STIL repression showed a significant reduction in nuclear β-catenin fraction, suggestive of inhibitive Wnt signaling (**Figure 6E**). There has been a recent report suggesting STIL mediated regulation of AKT protein in gastric cancer (21) and another one suggesting AKT mediated regulation of β-catenin in CRC (22). Thus, we hypothesized that STIL regulates β-catenin via regulating AKT in CRC. STIL silencing resulted in a significant decrease in AKT and p-AKT protein suggesting STIL mediated β-catenin regulation via AKT/pAKT regulation in CRC independent of Shh (**Figure 6C & D**). These results suggest that STIL functions in a hedgehog independent manner for regulation of β-catenin in CRC.

## 4 Discussion

STIL alteration has been implicated in lymphoblastic leukemia and microcephaly (23,24), however role of STIL in solid epithelial tumors stands least explored. Our report substantiates a novel role for STIL in regulation of tumor growth, drug resistance and stem cells in CRC. STIL plays an instrumental role in centriole division and thus has been very critical for cell cycle progression and proliferation(25). Being a regulator of cell proliferation its role in cancer becomes obvious and many tumor types have been reported to have enriched expression of STIL (4,26). In this study we found an over expression of STIL at mRNA and protein level in CRC tissues whereas, adjacent normal tissue showed a basal level expression (**Figure 1**). However, till date role of STIL in CRC growth and proliferation has not been studied. STIL being a critical factor in cell cycle machinery, its down regulation resulted in aberrant cell cycle and reduced cell proliferation in HT29 cancer cells (**Figure 3**). Further studies with STIL silenced xenograft showed a remarkable difference in tumor growth in NOD/SCID mice with lesser tumor volume and weight compared to control shRNA group. Ki-67 antigen, an established proliferation marker was also found to have less nuclear staining in STIL silenced tumor (**Figure 3**). Put together, these data showed a critical role of STIL in CRC proliferation and growth. Another aspect of STIL which has not been explored, is its role in hedgehog signaling and stem cell maintenance in cancer. CSC are being studied for development of future therapeutic interventions in CRC as they are known to be the fuel behind therapy failure and recurrence (27). Although CSCs are well known to be regulated by developmental signaling cascades, specific targets are lacking which can be exploited for development of advanced therapeutics. CD133 and CD44 surface molecules have been studied well in CRC for their enriched expression in CSC and responsible for drug resistance, recurrence and metastasis in CRC (28–30). Our study has shown that repression of STIL in HT-29 cells results in a significant reduction of CD133 & CD44 positive cells (**Figure 4**). Interestingly, STIL was found to regulate expression of CSC markers (**Figure 4**) and regulators such as OCT4 & NANOG at the transcriptional level both *in vitro* and *in vivo* (**Supplementary Figure 9**). These data suggest a stem cell regulatory function of STIL in CRC. However, when Shh signaling was inhibited by SANT1, we did not observe any significant reduction in CD133 and CD44 markers expression (**Figure 4**) along with OCT4 and NANOG genes (**Supplementary Figure 1**), which signs at an Shh independent regulation of CSC markers by STIL in CRC. Another hallmark feature of CSC and drug resistant cells is the ability to efflux drugs by enhancing ABC pumps expression (31). SP cells show high ABC transporter activities and CSC are reported to be enriched in SP cells in multiple gastrointestinal cancer (32). Hedgehog signaling has been reported to regulate expression of ABC transporters in ovarian epithelial cancer (33) and STIL being a positive regulator of GLI1 mediated hedgehog regulation was found to be significantly regulating side population in CRC cells (**Figure 5**). ABCG2 remains as one of the major efflux pump regulating side population as well as stem cells (34) and we observed down regulation of ABCG2 and partial reduction in ABCB1 expression upon STIL repression (**Figure 5**). Further, STIL silencing was found to repress expression of thymidylate synthase (**Figure 5**), a known target for 5-fu and a poor prognostic marker in CRC (35). To our concern we did not observe reduction of ABCG2 and thymidylate synthase protein upon Shh inhibition suggesting a critical role of STIL in regulation of drug resistant cells and genes in CRC independent of Shh signaling (**Figure 5**). Inhibition of STIL expression could sensitize HT29 cells for 5-fu treatment resulting in increased cell death (**Figure 5**), which has been supported by a previous study where STIL repression was shown to have a synergistic effect with DNA damaging agents in ovarian cancer (36). Conclusively, these data suggest that STIL could be a potent target for future therapeutics in CRC.

**Figure 5.**
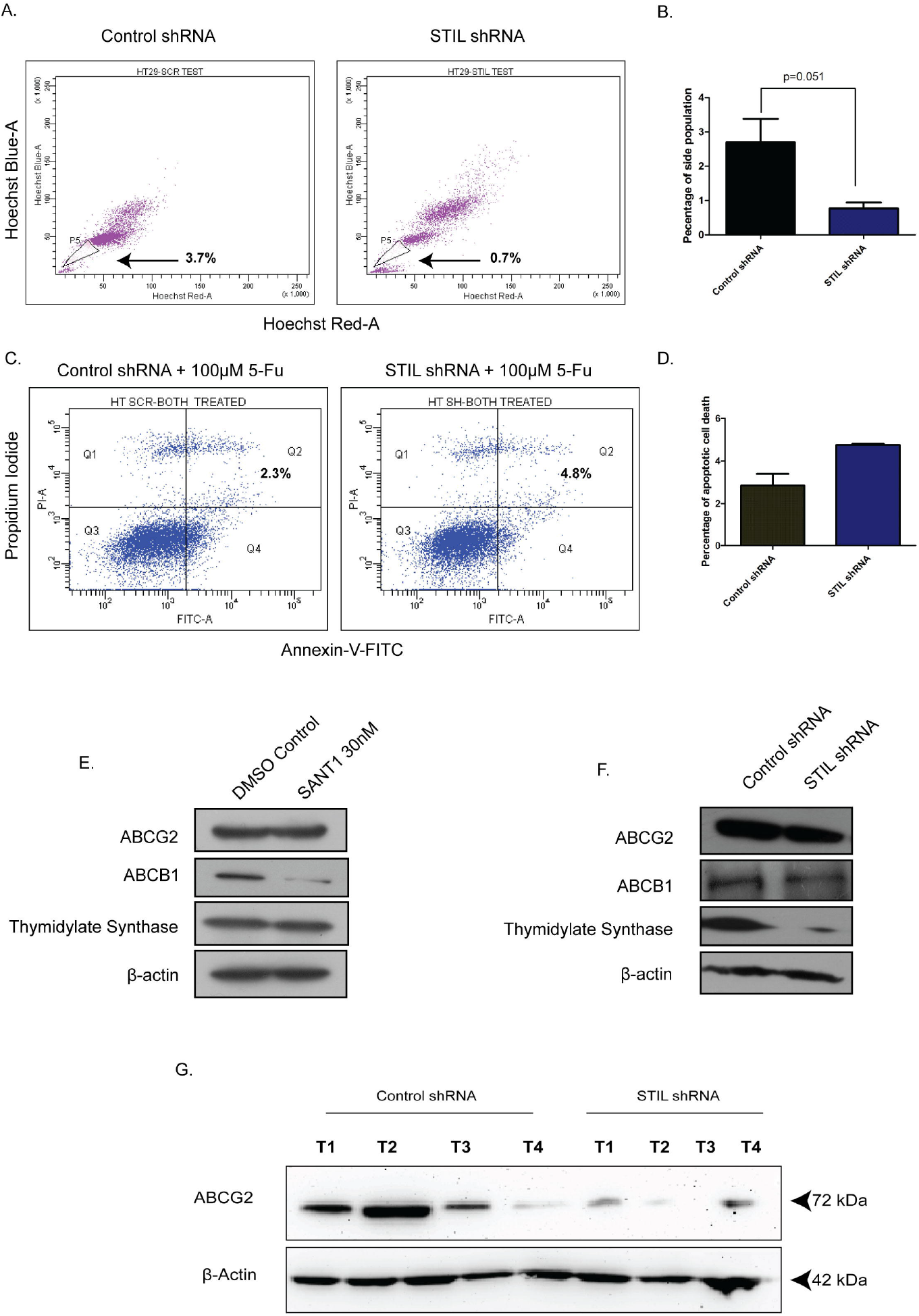
Effect of STIL silencing on drug effluxing cells and 5-FU therapy in CRC. (**A & B**) Dot plot showing side population cells and bar plot representing quantification of the same. (**C & D**) Annexin-V assay showing percentage of cell death upon 5-FU treatment. (**E & F**) Western blot showing expression of ABCG2,ABCB1 and thymidylate synthase protein expression upon SANT1 treatment and STIL silencing respectively. (**G**) Showing ABCG2 protein expression in control and STIL silenced tumor xenograft. Unpaired t-test results showing p value ≤0.05, is represented by *.

STIL has been established to be a positive regulator of Shh signaling and known to regulate GLI1 (37). Shh inhibition by SANT1 treatment showed reduced gene expression of its pathway component including GLI1/2, however STIL silencing was found to down regulate GLI1 and GLI2 but not other Shh components (**Figure 6**). These results advocate for an Shh independent regulatory function of STIL, which demands further studies. Shh and Wnt signaling has been studied for their antagonistic mode of regulation (20) and Wnt being the most critical aberrant signaling in CRC (38), we further looked into STIL mediated regulation of Wnt pathway. Interestingly, we found STIL to be regulating β-catenin protein, however Shh inhibition did not show similar result which further opens a new molecular axis of Wnt and Shh crosstalk mediated by STIL (**Figure. 6**). A recent study have delineated that STIL regulates PI3K/AKT in gastric cancer. Reiteratively, Akt is known to regulate β-catenin stabilization(39,40). These studies led us to explore an AKT mediated β-catenin regulation by STIL. One of the most interesting findings of our study remains to be STIL mediated regulation of β-catenin via AKT and p-AKT independent of Shh signaling (**Figure 6**). Eventually, our study revealed a STIL mediated cross talk between Wnt and Shh signaling in CRC, which opens new molecular insights on existing interplay between these developmental pathways during carcinogenesis and therapy failure in CRC. Despite few studies having shown the role of STIL in various cancers, its association with cancer prognosis stands elusive. Rectal tissue array analysis showed no association of STIL protein expression with invasive stages of tumor but was found to be highly expressed in early stages (**Figure 2 & Table 2**). This again may be supported by our scratch assay experiment where we have observed that STIL silencing have no significant effect on HCT116 migration and tumor xenograft also showed no tumor deposits in distant metastatic sites (**Supplementary Figure. 7**). Further overall survival (OS) and disease free survival (DFS) analysis from c-Bioportal revealed STIL over expression to be significantly associated with lower DFS in CRC but no association was found with OS (**Figure 1**). These observations show that STIL may not have a possible role in invasion but could be critical in early tumorigenesis and therapy resistance thus leading to lower DFS in CRC. However, more studies are warranted for elucidation of detailed molecular events mediated via STIL in CRC. Nonetheless our study suggest Shh and Wnt regulation by STIL to be a possible mechanism governing its role in CRC (**Figure 6 F**).

**Figure 6.**
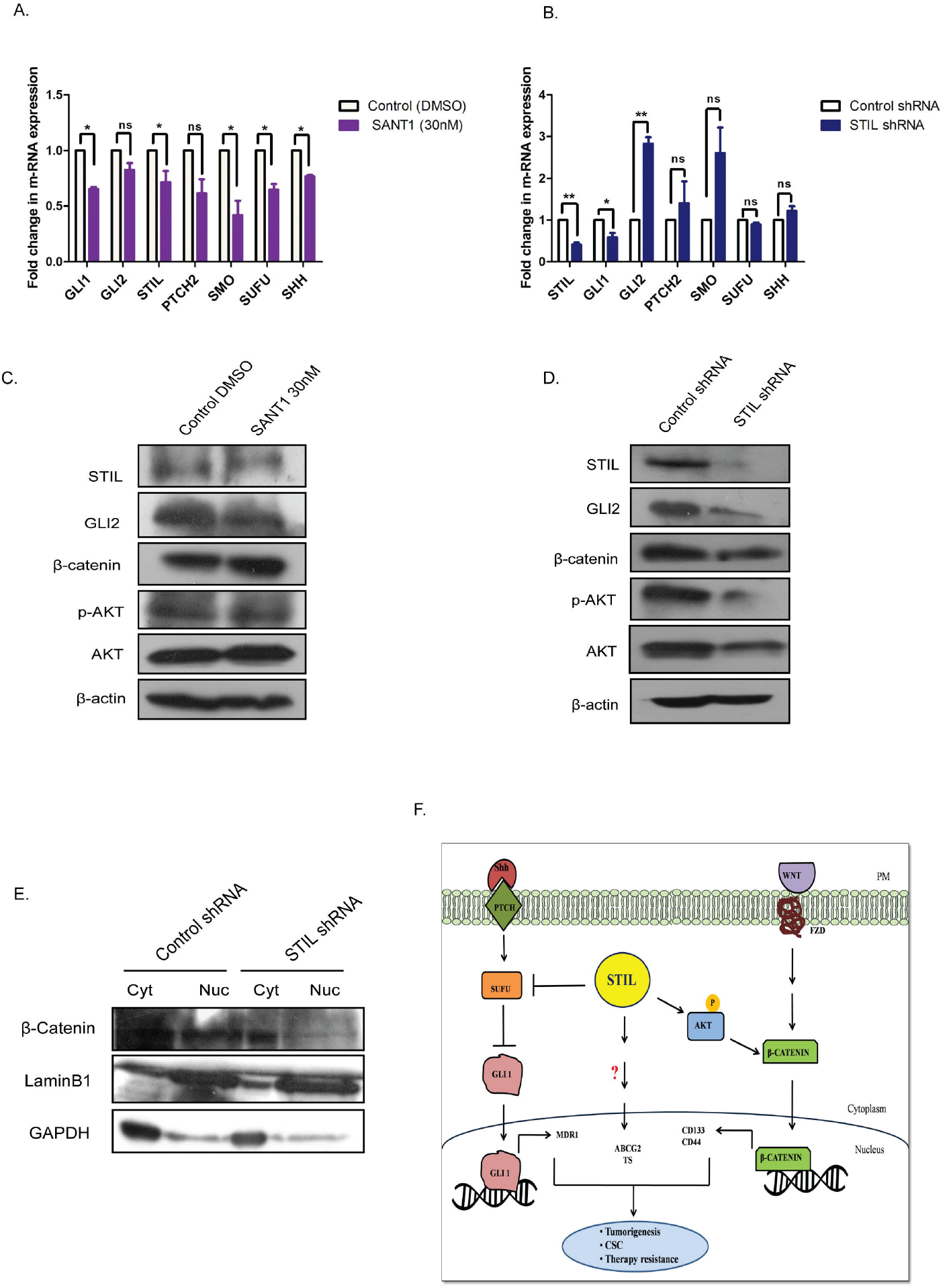
Effect of SANT1 treatment and STIL inhibition on Shh and Wnt signaling. (**A & B**) Bar graph showing expression of Shh signaling components upon SANT1 treatment and STIL silencing. (**C & D**) Western blot showing effect of SANT1 treatment and STIL silencing on GLI2, β-catenin, AKT and p-AKT protein expression. (**E**) Western blot showing β-catenin protein in cytoplasmic and nuclear fraction. (**F**) Schematic representation showing possible mechanistic regulation of multifaceted role by STIL in CRC. Unpaired t-test results showing p value ≥0.05, ≥0.01 are represented by *, ** respectively, ns-not significant.

In conclusion, this report stands as the first study demonstrating an oncogenic role of STIL in CRC. Our study showed the role of STIL oncogene regulating CSC characteristics along with drug resistance properties which had not been investigated till date in CRC. Further, STIL showed an Shh independent regulation of β-catenin, which we consider to be a novel finding, as regulation of Wnt by STIL opens a whole new prospective of research for Shh and Wnt cross talk in CRC. In addition, this study also sheds light on prognostic implications of STIL in CRC which ensue to be very critical and imperative from a clinical perspective. Association of STIL with poor disease free survival suggest its potency for a prognostic biomarker in CRC. Though, the mechanistic proof of STIL function in details remain elusive, our study urge that STIL could be an important oncogene with an inherent ability to regulate multiple aspects of cancer progression in CRC and it could be a warranted target for therapeutic intervention and prognostic marker in future.

## Availability of data and material

All the data analyzed and given in the manuscript are available from the corresponding author upon reasonable request.

## Authors’ Contribution

TP conceived, planned and carried out the experiments, lead in data analysis and manuscript writing. VK & SJ assisted in planning and execution of xenograft experiment. ESH & KR performed immunoblots and gave critical comments to improve work and also helped in article corrections. JVT helped with statistical analysis of IHC tissue array data. KC performed the surgery and helped in obtaining the human biopsies samples and their storage. AS helped in RNA isolation from biopsies and qPCR. AN conceived, designed the project and supervised its execution, manuscript proofreading and provided critical feedbacks in manuscript writing. All authors provided critical feedback and helped shape the research, analysis and manuscript.

## Acknowledgement

We would like to thank Dr. Arya, Dr. Vishnu & Dr Archana for their immense support in animal experiment. Dr. Debasree for HEK293T cells. Indu Ramachandran & Arya (Flow Cytometry facility, RGCB, Trivandrum) for her help and guidance in flow cytometry, Viji and Hima (Histology facility, RGCB, Trivandrum) for their assistance in histology work, Rekha for her help in biopsies collection.

## Ethical approval and consent to Participate

All human biopsies were collected at Regional cancer centre, Trivandrum, Kerala, India after approval of the Regional Cancer centre Ethical committee (HEC No.43/2011) and from donors that signed written informed consent. All the experiments on NOD-SCID mice were performed after approval from the Institute Animal Ethics Committee (IAEC/683/ASN/28).

