## Supplementary Figures for "STIL-a novel link in Shh and Wnt signaling, endowing oncogenic and stem like attributes to colorectal cancer"

**Supplementary Figure 7**

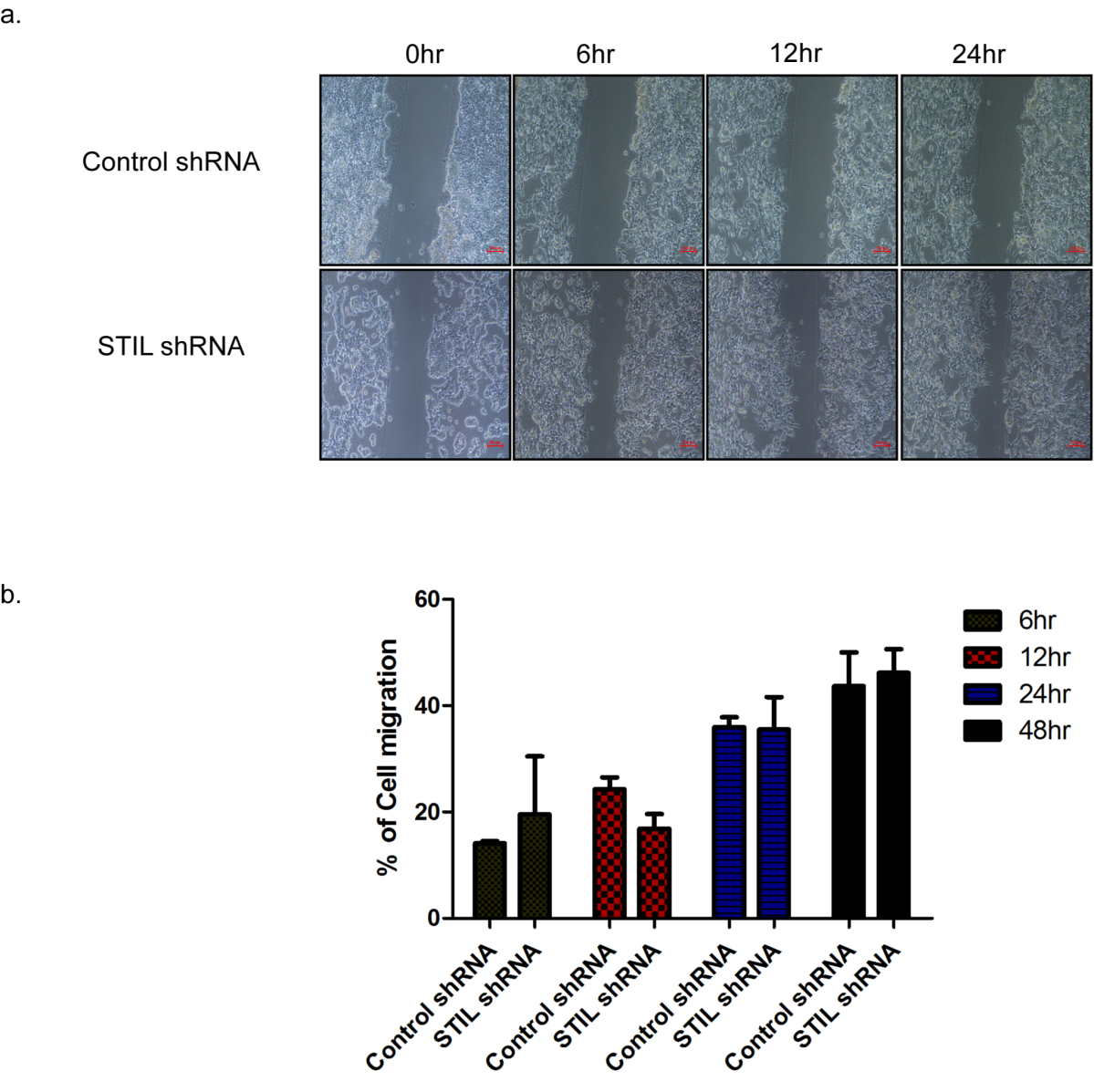

**Supplementary Figure 7 Showing effect of STIL silencing on cell migration**. A & B. Cell migration in control and STIL silenced HT29cells at different time points.

**Supplementary Figure 8**

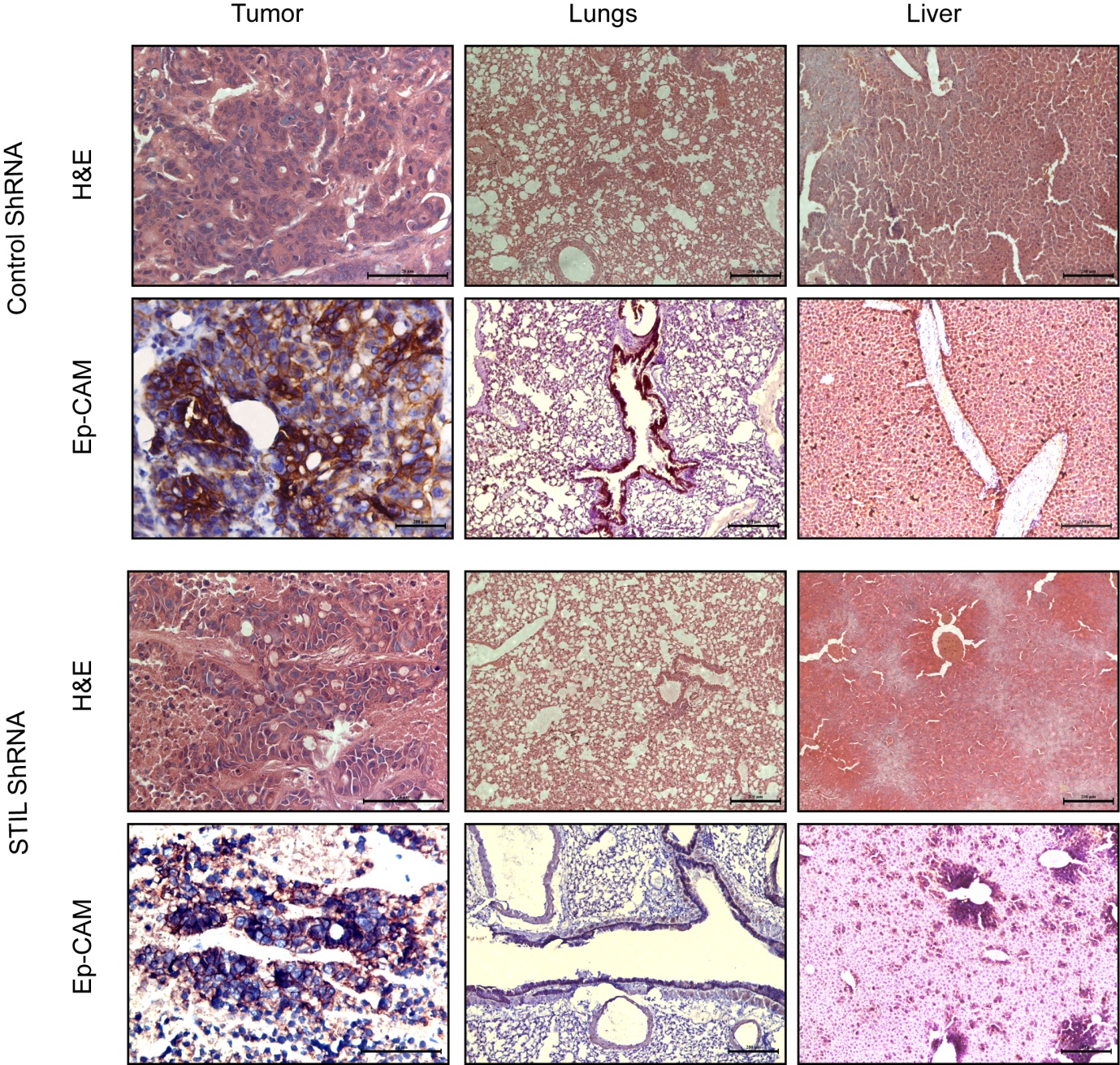

**Supplementary Figure 8 Showing Ep-CAM and H&E staining of control vs STIL silenced tumor xenograft, lungs and liver.**

**Supplementary Figure 9**

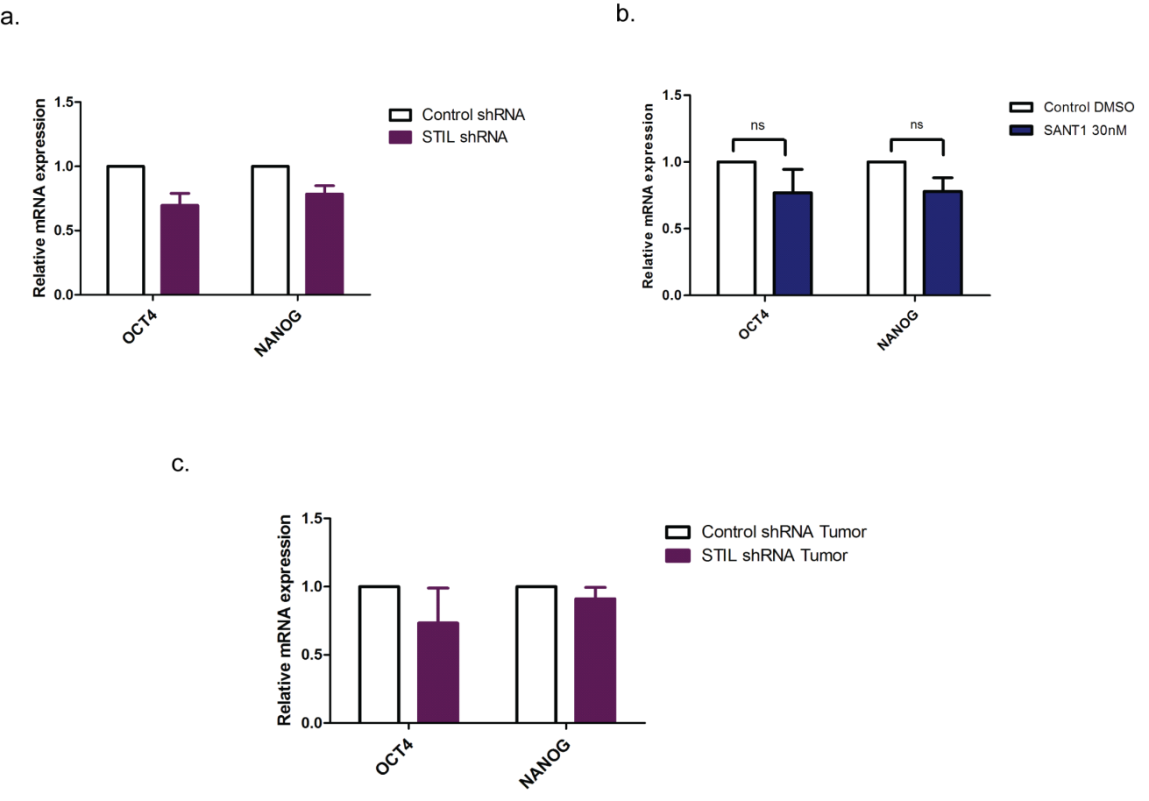

**Supplementary Figure 9** Realtime qPCR data showing expression of OCT4 & NANOG in CRC upon STIL silencing and SANT1 treatment.

**Supplementary Figure 10**

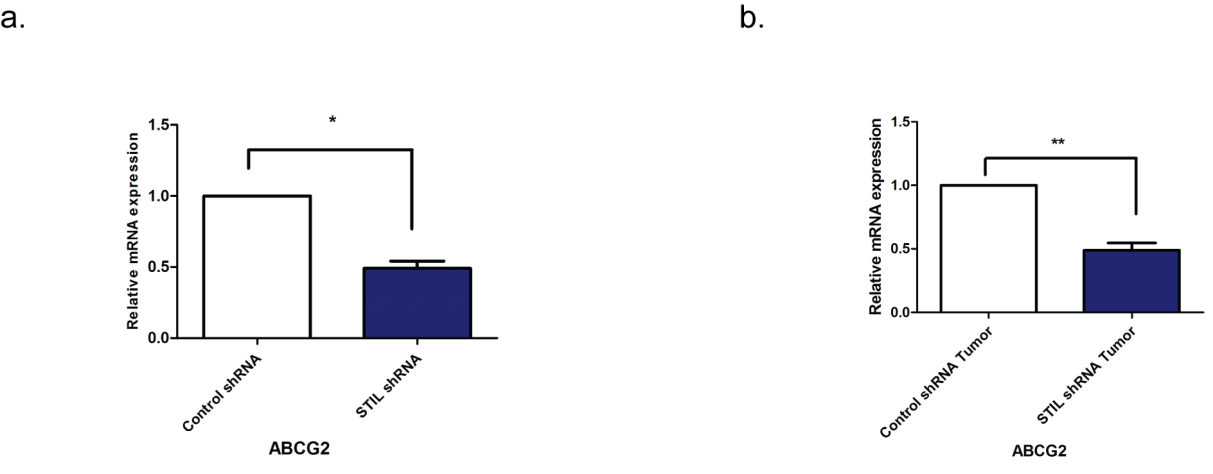

**Supplementary Figure 10 Showing effect of STIL silencing on expression of ABCG2 gene in CRC.** A. ABCG2 expression in control and STIL silenced HT29 cells. B. ABCG2 expression in control and STIL silenced Xenograft tumors.

**Supplementary Figure 11**

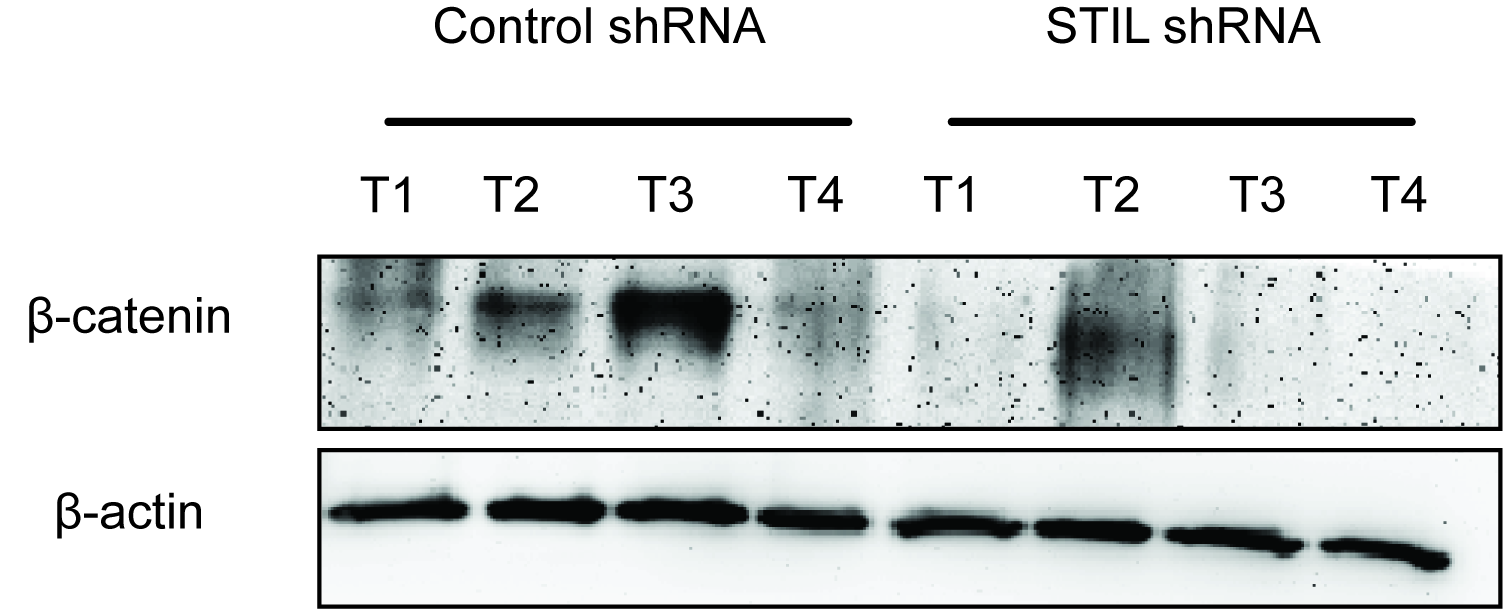

**Supplementary Figures 11 Showing expression of β-catenin protein upon STIL silencing.** Expression of β-catenin in control and STIL silenced tumor xenograft. (T- Tumor, 1-4 represents different mice)
