## Supplementary Tables for "STIL-a novel link in Shh and Wnt signaling, endowing oncogenic and stem like attributes to colorectal cancer"

**Supplementary Table 1**

**Primer List :**

A total of nine genes were evaluated for gene expression using real-time q-PCR in normal, tumor and distal biopsies. Details of both forward and reverse primers sequences are given in this table.

| **Sr. No.** | **Genes Name** | **Primer Sequence 5' to 3' -** |
| --- | --- | --- |
| 1 | CD133-F | GCATTGGCATCTTCTATGGTT |
| 2 | CD133-R | CGCCTTGTCCTTGGTAGTGT |
| 3 | CD44-F | AGAAGGTGTGGGCAGAAGAA |
| 4 | CD44-R | AAATGCACCATTTCCTGAGA |
| 5 | OCT4-F | GGGAGATTGATAACTGGTGTGTT |
| 6 | OCT4-R | GTGTATATCCCAGGGTGATCCTC |
| 7 | NANOG-F | CAGAAGGCCTCAGCACCTAC |
| 8 | NANOG-R | ATTGGAAGGTTCCCAGTCGG |
| 9 | STIL -F | GGATGTCCAGCCCTGTACTGT |
| 10 | STIL-R | GCCACAGGCGATGGTTCAAT |
| 11 | PTCH2-F | CCAGCCCTCGCTTCTTGATT |
| 12 | PTCH2-R | GCCTATGCCTTCCTCAGGTG |
| 13 | SUFU-F | GCCTGAGTGATCTCTATGGTGA |
| 14 | SUFU-R | TCTCTCTTCAGACGAAAGGTCAA |
| 15 | SHH-F | GGCCAAAGCGTTCAACTTGT |
| 16 | SHH-R | TTGCCCAGGCTAACGTGTC |
| 17 | SMO-F | CCTGTGGCAACTCCAGGTATG |
| 18 | SMO-R | GATCCTCTGGGGCCTTTCCT |
| 19 | GLI1-F | GAGCCAGAAGTTGGGACCTC |
| 20 | GLI1-R | CCTCGCTCCATAAGGCTCAG |
| 21 | GLI2-F | TGCAACGTCCACCCACTTTA |
| 22 | GLI2-R | GGATACCAGGCATGCACACT |
| 17 | L19 -F | GCGGAGAGGGTACAGCCAAT |
| 18 | L19 -R | GCAGCCGGCGCAAA |

**Supplementary Table 2**

**shRNA Details:**

| **Gene Name** | **shRNA sequence** | **REFSEQ_ID** | **Company** |
| --- | --- | --- | --- |
| **STIL** | CCGGGTCTGGAATTACACATATCTACTCGAGTAGATATGTGTAATTCCAGACTTTTT | NM_001048166.1,NM_003035.2 | Dr G. Subba Rao  MCB,IISC,  Bangalore |

**Supplementary Table 3**

Table shows the details of antibodies used in Flow cytometry, Western blot , Immunofluroscent imaging and Immunohistochemsitry.

| **Antibody Name** | **Company name and Catalog No.** |
| --- | --- |
| Anti-Ep-CAM | Abcam,ab 71916, USA |
| Anti-CD44 | Abcam,ab157107, USA |
| Anti-OCT4 | Abcam,ab18976, USA |
| Anti-β Catenin | CST,8480S, USA |
| Anti-CD133 | CST,#86781,USA |
| CD133 (AC133)-PE | Miltenyi,130-098-826,Germany |
| CD44-FITC | Miltenyi, 130-098-210,Germany |
| CD326-VioBlue | Miltenyi, 130-098-092,Germany |
| Anti-Ki67 | Abcam,ab16667,USA |
| Anti-STIL | Abcam,ab89314,USA (Dilution WB-1:1000) |
| Anti-β-catenin | CST,8480,USA (Dilution WB-1:1000) |
| Anti-β-actin | Sigma-aldrich,A5316,USA (Dilution WB-1:5000) |
| Anti-ABCG2 | Abcam ,108312,USA (Dilution WB-1:1000) |
| Anti-Thymidylate synthase | CST,9045,USA (Dilution WB-1:1000) |
| Anti-GLI2 | Abcam ab26056 (Dilution WB-1:500) |
| Anti cMyc | Abcam ab39688 (Dilution WB-1:1000) |
| Anti-MDR1 | Abcam ab170904 (Dilution WB-1:1000) |
| Anti-pAKT(Thr 308) | CST 244F9 (Dilution WB-1:2000) |
| Anti-AKT | CST 9272 (Dilution WB-1:2000) |

**Supplementary Table 4**

Rectal carcinoma tissue array cases demographics.

| **Patients characteristics** | | |
| --- | --- | --- |
| **Total number of patients** |  | 75 |
| **Median Age** |  | 55 Years |
| **Sex** |  |  |
|  | Women | 36 (48%) |
|  | Men | 39(52%) |
| **Pathology of diagnosis** |  |  |
|  | Normal | 5 (6.67%) |
|  | Adenocarcinoma | 54 (72%) |
|  | Mucinous Adenocarcinoma | 8 (10.67%) |
|  | Signet-ring cell carcinoma | 5 (6.67%) |
|  | Others | 3 (4%) |
| **Depth of Invasion** | T1 | 5 (7.14 %) |
|  | T2 | 25 (35.71%) |
|  | T3 | 28 (40%) |
|  | T4 | 12 (17.14%) |
| **Lymph node metastasis** |  |  |
|  | Negative | 63 (90%) |
|  | Positive | 7 (10%) |
| **Distant metastsis** |  |  |
|  | Negative | 70 (100%) |
|  | Positive | 0 |
| **Histologic type** |  |  |
|  | Grade 1 | 18 (29.5%) |
|  | Grade 2 | 33 (54.1%) |
|  | Grade 3 | 10 (16.39%) |
|  | Unknown | 9 (12.8%) |
| **Site of tumor** |  |  |
|  | Rectum | 75(100%) |

**Supplementary Table 5**

**Table shows buffer composition.**

| CE Buffer(pH 7.6) | |
| --- | --- |
| 10 mM HEPES(4-(2-hydroxyethyl)-1-piperazineethanesulfonic acid) | Gibco, Invitrogen |
| 60 mM KCl | EMPARTA, Merck Lifescience |
| 1 mM EDTA | Sigma-Aldrich, USA |
| 0.075% (v/v) NP-40 | Sigma-Aldrich, USA |
| 1mM Dithiothreitol(DTT) | Sigma-Aldrich, USA |
| 1 mM Phenylmethane sulfonyl fluoride(PMSF) | Merck |
| NE Buffer (pH 8.0) | |
| 20 mM Tris-Cl | EMPARTA, Merck Lifescience |
| 420 mM NaCl | EMPARTA, Merck Lifescience |
| 1.5 mM MgCl2 | Sigma-Aldrich, USA |
| 0.2 mM EDTA | Sigma-Aldrich, USA |
| 1mM PMSF | Merck |
| 25% (v/v) Glycerol | Sigma-Aldrich, USA |
